# Single-molecule tracking of Nanog and Oct4 in mouse embryonic stem cells

**DOI:** 10.1101/2022.08.08.503148

**Authors:** Kazuko Okamoto, Hideaki Fujita, Yasushi Okada, Soya Shinkai, Shuichi Onami, Kuniya Abe, Kenta Fujimoto, Tomonobu M Watanabe

## Abstract

Nanog and Oct4 are core transcription factors in a gene regulatory network that regulate hundreds of target genes for pluripotency maintenance in mouse embryonic stem cells. To understand their function as a gatekeeper or a pioneer factor at the molecular scale, we quantified the residence time on target loci, fluctuation at the loci, and interaction clustering by visualizing single molecules of Nanog and Oct4 in a living nucleus during the pluripotency loss. Interestingly, Nanog interacted longer with its target loci in the lower Nanog expression state or at the onset of differentiation, indicating the possibility of a new feedback mechanism to maintain Nanog expression. The interaction time of Nanog and Oct4 corresponded to their fluctuation and interaction clustering, which depended on their expression or differentiation state, respectively. The DNA viscoelasticity near the Oct4 target locus remained flexible during the differentiation, reflecting its role as a pioneer factor. Based on these results, we propose a new feedback mechanism for pluripotency maintenance in which Nanog function is prolonged, corresponding to chromatin condensation, and Oct4 reopens the condensation.

## Introduction

The starting cells of differentiation are the embryonic stem cells (ESCs), which can self-renew and differentiate to most types of cells, termed pluripotency (Beddington & Robertson 1989). Cell fate determination in differentiation is driven by a dynamic interplay of biological reactions in the nucleus where the expression of each gene is switched on or off by the binding of the transcription factors (TFs) to their target loci, resulting in patterned expressions of regulatory genes. The binding of TFs is regulated by chromatin condensation–relaxation controlled by histone and/or DNA modifications (Lambert *et al*., 2018; Peñalosa-Ruiz *et al*., 2019). Though ESCs are thought to irreversibly lose pluripotency during differentiation to other somatic cells, Takahashi and Yamanaka demonstrated that pluripotency could be artificially induced/resurrected in somatic cells by exogenously expressing only four specific TFs (Takahashi & Yamanaka, 2006). This experimental fact suggests a cyclic nature of the interplay of the transcription process, epigenetic modification, and chromatin condensation–relaxation. TFs that facilitate effective transcription of their target genes to regulate hundreds of downstream genes for pluripotency induction and maintenance are called core TFs. A working model of how the core TFs interact in the nucleus for pluripotency maintenance contributes to elucidating the mechanism underlying differentiation and reprogramming and effectively enhances the production and quality control of induced pluripotent stem (iPS) cells. Many researchers have tackled this issue with various advanced technologies in distinct research fields.

Herein, we focused on two core TFs, Nanog and Oct4, which co-work to stabilize ESCs in the pluripotent state; Nanog buffers the differentiation activity mxediated by Oct4 (Loh *et al*., 2006; Liang *et al*., 2008). They induce their own expression as well as mutually activate others; thus, resulting to a positive feedback circuit (Pan & Thomson, 2007). For Nanog, by applying molecular expression noise to the feedback circuit, ESCs stochastically fluctuate between two stable states: high and low Nanog expressions (Herberg & Roeder, 2015; Marucci, 2017). It has been suggested that the gate to differentiation can be opened only when it is in a low expression state. Hence, Nanog is called a “gatekeeper” (Hyslop *et al*., 2005). Generally, chromatin condenses during differentiation from an open structure that exposes interacting sites for TF binding to a closed structure to prevent them (Gaspar-Maia *et al*., 2011; Apostolou & Hochedlinger, 2013). Oct4 is one of the pioneer factors responsible for reopening/remodelling closed chromatin (Soufi *et al*., 2012; Iwafuchi-Doi & Zaret, 2014; Xiong *et al*., 2022). Numerous studies support these hypotheses. Along with these, computational experiments interpolate the pieces of static evidence. Nevertheless, the dynamic observation of functioning proteins on-site is required to prove these hypotheses, as to the final evidence, and/or propose a new hypothesis.

Single-molecule tracking (SMT) based on fluorescence microscopy is the most appropriate tool for investigating the dynamic function of individual protein molecules of interest in living cells (Liu & Tjian, 2018; Shao *et al*., 2018; Lionnet & Wu, 2021). A protein of interest is conjugated with a fluorescent dye one-to-one, and even the weak fluorescence emitted from the single dye molecule can be detected if the background fluorescence is sufficiently low. However, the background fluorescence in conventional fluorescence microscopy veils a weak fluorescent signal. Total internal reflection fluorescence microscopy (TIRFM) improves the signal-to-noise ratio (S/N) by 2,000 times through selective illumination that uses evanescent light generated when the incident light is totally reflected at the interface boundary (Funatsu *et al*., 1995). Thereby, the fluorescence-coupled protein is visualized as a simple fluorescent spot. TIRFM is limited to observing proteins on or near the plasma membrane attached to a glass surface since the evanescent field is localized on the medium–coverslip interface (Sako & Uyemura, 2002). To observe single molecules inside the nucleus, a thinned sheet-formed light illumination approach has been developed, which achieved a sufficient S/N to detect signals emitted from single dye molecules, although its improvement of the S/N is not as effective as TIRFM (Tokunaga *et al*., 2008; Gebhardt *et al*., 2013; Izeddin *et al*., 2014). Nevertheless, the in-nucleus SMT has been achieved to quantitatively investigate the single-molecular behavioral characteristics of TFs by obtaining kinetic parameters, such as residence time on target loci, fluctuation at the loci, and interaction clustering (Liu & Tjian, 2018; Shao *et al*., 2018; Lionnet & Wu, 2021).

The distribution of fluorescence emitted from a single individual fluorophore onto the detector surface, called a point spread function (PSF), approximately follows a Gaussian distribution (Thompson *et al*., 2002). The PSF distribution can be used as a weight to calculate the centre position, intensity, and radial width of a fluorophore. SMT requires the identification and collection of these parameters of many fluorophores as possible in all acquired images and applying them to appropriate analyses for each object (Liu & Tjian, 2018; Shao *et al*., 2018; Lionnet & Wu, 2021). Since it is time-consuming and labour-intensive to identify each fluorescent spot one-by-one by human eyes, some methods to automatize this task have been proposed in previous studies (Mashanov & Molloy, 2007; Wilson *et al*., 2016; Yasui *et al*., 2018). These automatic identifications work well in cases with sufficiently high S/N values. However, in the thinned sheet-formed light illumination, diffusive scattering and refraction due to intracellular microstructure by irradiating a cell from the side cause background speckles, which could be misidentified when using low intense dye, such as genetically-encoded fluorescent proteins, or when the dye concentration is high.

Self-labeling proteins, such as Halo-tag (England *et al*., 2015), CLIP-tag (Gautier *et al*., 2008), and SNAP-tag (Keppler *et al*., 2003; Keppler *et al*., 2004), can address this problem, and have been used in SMT of TFs (Chen *et al*., 2014; Liu *et al*., 2014; Xie *et al*., 2017; Piccolo *et al*., 2019; Gómez-García *et al*., 2021). The self-labeling protein does not emit fluorescence, but can fluorescently label a protein of interest via a ligand conjugated with stable and intense fluorescent dye loaded into a cell. Therefore, the users can adjust the labeled protein concentration appropriate for their SMT condition and analysis object, thereby effectively reducing the misidentification and elongating the number of tracking frames. However, the self-labeling protein does not apply to this study. For Nanog or Oct4, which activates its own expression, it is important to investigate the relationship between the single-molecule behavioral characteristics and its fluorescence intensity, a measure of the expression level, in the same cell. Due to cellular heterogeneity in dye uptake, it is almost practically impossible to maintain a constant labeling efficiency across the cells, resulting in a mismatch between the fluorescence intensity and the expression. Genetically-encoded fluorescent proteins are more advantageous than self-labeling proteins depending on the case. This dilemma is a common limitation during single-molecule observation of TF working in living cell nuclei. New methods to overcome the lower S/N of in-nucleus SMT with genetically-encoded fluorescent proteins are still needed.

Herein, we performed a direct on-site observation of single molecules of Nanog and Oct4 fused with a fluorescent protein in a living cell nucleus during the pluripotency loss in mouse ESCs (mESCs) with careful construction of a method of in-nucleus single-molecule analysis. The quantitative analysis results highlighted the differences between Nanog and Oct4 in single-molecule dynamics. Also, the results implied the roles of Nanog as a gatekeeper and Oct4 as a pioneer factor, respectively, biophysically proving the current hypotheses. Furthermore, according to these results, we propose a new feedback mechanism for pluripotency maintenance by co-functioning Nanog and Oct4. Hence, this study demonstrates the veracity of the current hypothesis and provides novel insights into pluripotency maintenance based on single-molecule quantification.

## Results

### Single-molecule observation of Nanog or Oct4 fused with EGFP in mESCs

Endogenous Nanog or Oct4 in mESCs were labeled with a variant of green fluorescent protein derived from *Aequorea victoria* (Shimomura *et al*., 1962; Tsien, 1998), enhanced GFP (EGFP) (Heim *et al*., 1995) (Fig. 1A, *upper*). Reasons for selecting EGFP among GFP variants are described in Supplemental Text (I) and Figs. S1 and S2. The nucleus region in an mESC expressing Nanog- or Oct4-EGFP was selectively illuminated with a sheet-formed laser using highly inclined and laminated optical sheet microscopy (HILOM) (Tokunaga *et al.*, 2008; Izeddin *et al*., 2014) (Fig. 1B). Nanog- or Oct4-EGFP molecules in the nucleus, indiscriminable under an epi-fluorescent microscope (Fig. 1C, *second left*), were unclearly visualized even under the HILOM because the EGFP molecule concentration was extremely high. Once all EGFP within the irradiated area in the HILOM was photobleached, only molecules newly coming into the irradiated area were discriminably visualized as fluorescent spots (Fig. 1C, *second right*). Because the diffusion of unbound proteins in the nucleus was too fast to be detected during the set exposure time (50 ms), a visible fluorescent spot corresponded to a Nanog- or Oct4-EGFP molecule stably interacting with its target loci on DNA. The duration of the interaction was defined from the appearance to the disappearance of a fluorescent spot on a certain site as dwell (Fig. 1D and supplemental Video S1). The repeated dwell events at the same location indicate that Nanog- or Oct4-EGFP molecules bind to the same locus (Fig. 1D, *yellow lines*).

**Figure 1.**
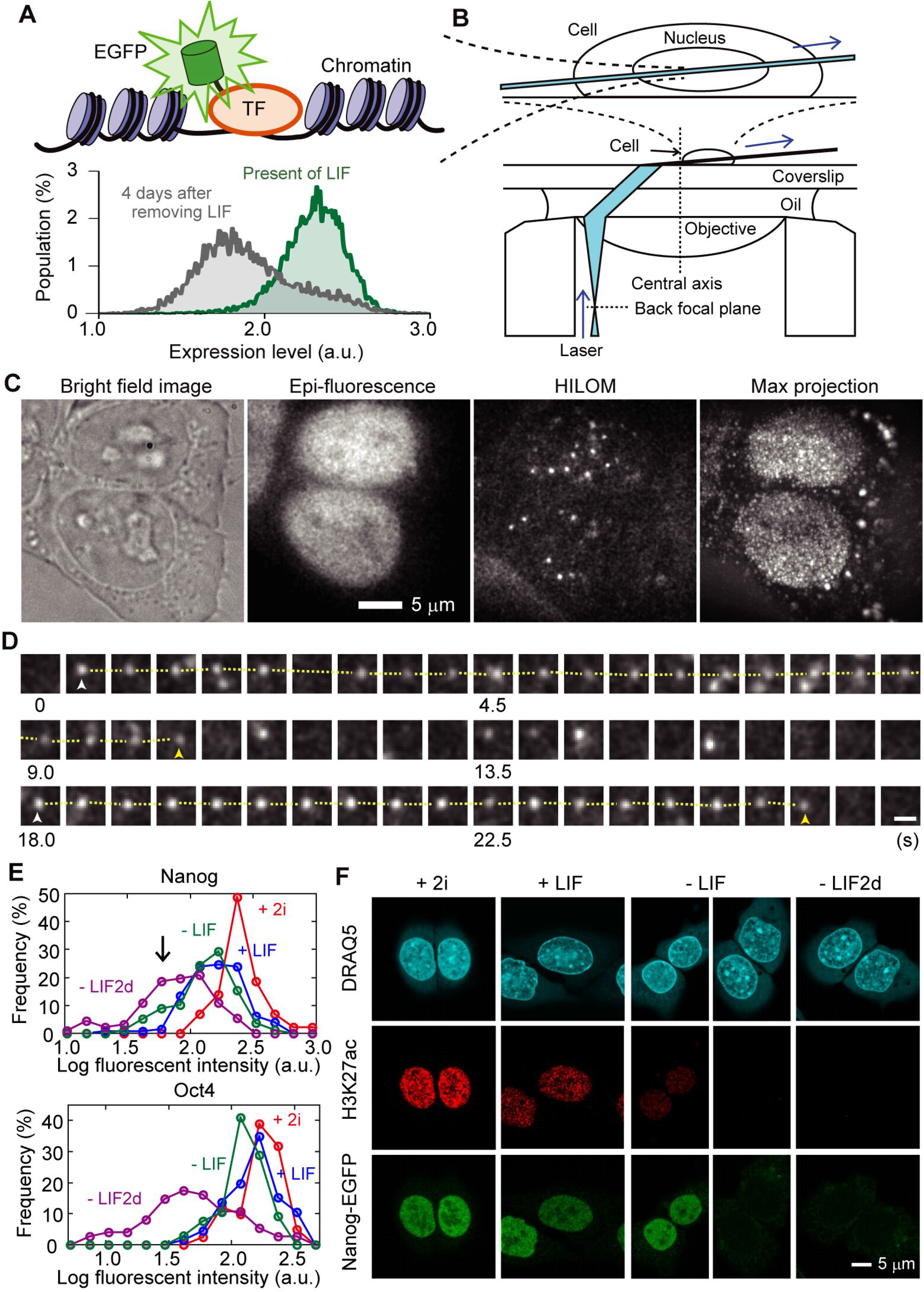
Single-molecule tracking of Nanog or Oct4 fused with EGFP in mESCs. (**A**) Cartoon of EGFP fused Nanog or Oct4, and a flow cytometry result of Nanog-EGFP expressing mESCs in the presence of LIF (*green*) and 4 days after removing LIF (*grey*). TF, Nanog or Oct4. (**B**) Schematic drawing of the present observation with HILOM. To observe a cell at 2–3 μm above the cell surface attached on a glass, the cell was placed slightly lateral to the centre of the field of view. (**C**) Representative image of mESC expressing Nanog-EGFP in +LIF. Bright field, epifluorescence, HILOM, and max projection image of 2,000 frames are shown. (**D**) Typical time course of a Nanog-EGFP molecule at 450 ms interval in +LIF. Yellow dotted lines indicate the dwell on the same site. White and yellow arrow heads indicate the appearance and disappearance of Nanog-EGFP, respectively. The scale bar is 1 μm. It is speculated that the chromatin near the loci where Oct4 is interacting opened when its expression decreased. (**E**) Histogram of average fluorescent intensity in an mESC expressing Nanog-EGFP (*upper*) or Oct4-EGFP (*lower*) at various differentiating condition defined as following; the presence of both LIF and 2i (+2i, *red*) as the initial condition, one day after removing 2i (+LIF, *blue*), further removal of LIF for one day (-LIF, *green*), and after one another day (-LIF2d, *magenta*). (**F**) Confocal images of DRAQ5 staining, showing double stranded DNA (*top*), immunostaining of H3K27ac (*middle*), and Nanog-EGFP (*bottom*), along with the progression of differentiation (+2i, +LIF, −LIF, and −LIF2d).

LIF enhances the Nanog and Oct4 expression, which is necessary to maintain stemness (Matsuda *et al*., 1999; Hirai *et al*., 2011). Additionally, MAPK/ERK and GSK3 inhibitors, named 2i, stabilize the pluripotency of mESCs to differentiate into germlines (Wray *et al*., 2010). The differentiating conditions were defined by the presence or absence of LIF and/or 2i, and the elapsed days since the removal of both (see Methods). The downregulation of Nanog- or Oct4-EGFP expression due to the LIF removal could be easily confirmed by observing the decrease in the fluorescent cells with fluorescence flow cytometry (Fig. 1A, *lower*). However, while the cells in a flow cytometer are unbound, the cells are bound on the glass surface in the HILOM. Cell adhesion to the substrate affects the Nanog and Oct4 expression in mESCs (David *et al*., 2019). Therefore, we confirmed the downregulation in our mESC lines under an epi-fluorescent illumination before performing SMT (Fig. 1C, *second left*). In the presence of both LIF and 2i (+2i), all Nanog-EGFP expressing mESCs exhibited high fluorescence (Fig. 1E, *upper, red*). One day after removing 2i (+LIF), the fluorescence intensity decreased, but the EGFP fluorescence remained detectable in almost all mESCs (Fig. 1E, *upper, blue*). Further removal of LIF for one day (-LIF) results in fluorescence loss in some mESCs (Fig. 1E *upper, green, arrowhead*). After one another day (-LIF2d), the cells turned nonfluorescent or too dark to be detected (Fig. 1E, *upper, magenta*). In the case of Oct4-EGFP expressing mESCs, the fluorescence was maintained until the first day after removing LIF and was almost lost by the second day (Fig. 1E, *lower*).

Next, we confirmed the chromatin condensation and acetylation of histone H3 at lysine 27 (H3K27ac) that depend on the differentiating condition using confocal fluorescence microscopy (Fig. 1F). Bright foci are interspersed in the DNA staining image (Fig. 1F, *top*), the fluorescence signal decreases in the H3K27ac immunostaining (Fig. 1F, *middle*), and Nanog-EGFP-negative mESCs are present (Fig. 1F, *bottom*), which all depend on the differentiation condition. EGFP molecules were undetected in EGFP-negative mESCs, even when using the HILOM. Inevitably, we selectively observed Nanog- or Oct4-EGFP expressing mESCs. Thus, we only observed mESCs just before the pluripotency loss.

### Optimization of autoidentification protocol of single molecules

To estimate the centre position (*x*_0_ [pixels], *y*_0_ [pixels]), intensity (*I* [a.u.]), and radial width (*σ* [pixels]) of a single fluorescent spot, a method was selected based on Gaussian fitting in which the parameters were obtained by fitting an image of a single fluorophore within a circle-formed region of interest (ROI) of nine pixels in diameter. This selection was made because the Gaussian fitting has the highest robustness at low S/N data compared to other methods, such as cross-correlation, sum-absolute difference, or simple centroid calculation (Thompson *et al.*, 2002). Additionally, we assumed a tilted planar background in the limited area within the ROI in the Gaussian fitting approximation (Fig. 2A and see Methods) (Ichimura *et al*., 2014). As truly identified molecules, we collected data on more than 8,645 single fluorescent spots that were individually identified by the human eye (hereinafter, “manual picking”) (Fig. 2B, *black*). Here the ROI where the iterative calculation solution for the Gaussian fitting converges was comprehensively stored by scanning within the nucleus region in a collection. The *I–σ* correlation plot of all calculated solutions exhibits a wider distribution than that of the manual picking (Fig. 2B, *green*), indicating that the solution for the Gaussian fitting might converge even if no fluorescent molecule exists. Therefore, the threshold *I* (*Th*_(*I*)_ [a. u.]) and/or the threshold *σ* (*Th*_(*σ*)_ [pixels]) are generally used to distinguish between ROIs with and without a fluorescent spot (Fig. 2BC, *broken red lines*) (Yasui *et al*., 2018).

**Figure 2.**
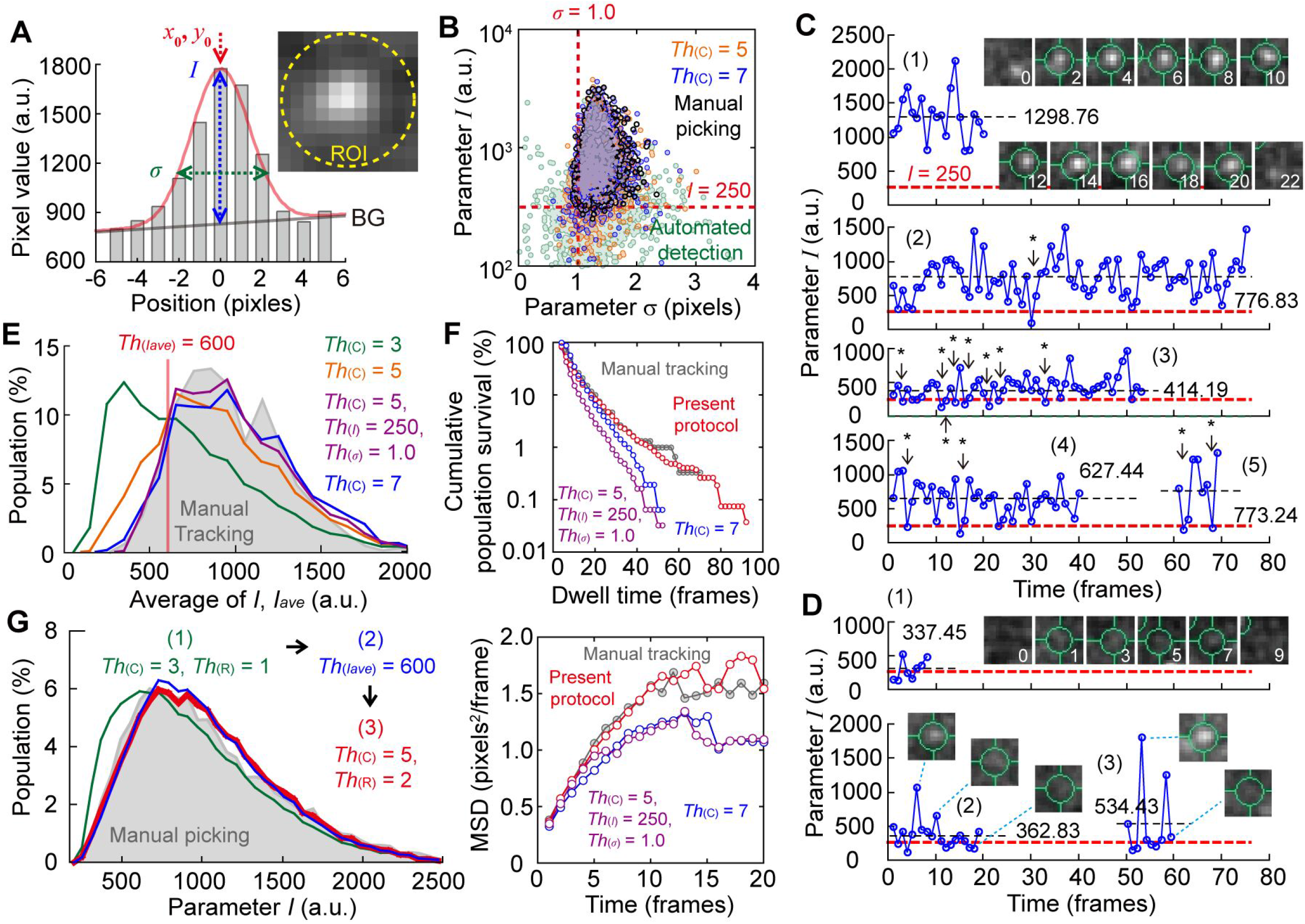
Optimization of auto-identification protocol of single individual molecules. (**A**) Typical cross-section of a fluorescent spot of Nanog-EGFP in an mESC and explanation of the fitting parameters. Red line is the fitting result with a Gaussian function with assuming the planar background (*BG*). The inset is a fluorescent image of single Nanog-EGFP molecule. Diameter of a circle ROI was 9 pixels (*yellow broken line*). (**B**) *I–σ* plot of the results of Gaussian fitting for Nanog-EGFPs in a nucleus by manual picking (*black*), ones by the automatized identification (*green*), and ones screened with thresholds of *Th*_(*C*)_ = 5 (*orange*) and *Th*_(*C*)_ = 7 (*blue*). (**C**) Four typical examples of time courses of *I* in a trajectory of true single molecules. (**D**) Four typical examples of time course of *I* in a trajectory including false positives. Insets in C and D are the time course of the ROI (*cyan circle*) in which *I* was estimated. Values in C and D indicate the average value of *I* in a trajectory, *I*_ave_. Red broken lines indicate *I* = 250, respectively. Asterisks indicate blinking points when *Th*_(*I*)_ = 250. (**E**) Histograms of *I*_ave_ of the result screened with thresholds of *Th*_(*I*)_ = 0, *Th*_(*σ*)_ = 0, *Th*_(*R*)_ = 1 and *Th*_(*C*)_ = 3 (*green*), = 5 (*orange*), or = 7 (*blue*). The one obtained by manual picking was superimposed in *grey. Magenta* is one with thresholds of *Th*_(*I*)_ = 250, *Th*_(*σ*)_ = 1.0, *Th*_(*R*)_ = 1 and *Th*_(*C*)_ = 5. (**F**) Cumulative histogram of dwell time (*upper*) and MSD plot (*lower*) using single-molecule data of Nanog-EGFP screened by manual tracking (*grey*) and with thresholds of *Th*_(*R*)_ = 1, *Th*_(*C*)_ = 5, *Th*_(*I*)_ = 250 and *Th*_(*σ*)_ = 1 (*magenta*) or *Th*_(*R*)_ = 1, *Th*_(*C*)_ = 7, *Th*_(*I*)_ = 0 and *Th*_(*σ*)_ = 0 (*blue*), and the present protocol (*red*). (**G**) Histograms of *I* of the result screened by the present protocol: *green,* the first screening with *Th*_(*C*)_ = 3 and *Th*_(*R*)_ = 1; *blue,* the second screening with *Th*_(*Iave*)_ = 600; *red*, the final screening with *Th*_(*C*)_ = 5 and *Th*_(*R*)_ = 2. The one obtained by manual picking was superimposed in *grey.*

Since TFs bind and remain in a locus, trajectories of time development after a spot appearance within a limited range (*Th*_(*R*)_ [pixels]) could be extracted using the threshold for the continuity of the fluorophore (*Th*_(*C*)_ [frames]) (Fig. 2CD), resulting in screening out misidentification, even without thresholds using *Th*_(*I*)_ nor *Th*_(*σ*)_ (Fig. 2B, *orange and blue*) (Mashanov & Molloy, 2007; Wilson *et al*., 2016). However, the collected trajectories with the thresholds of *Th*_(*R*)_ and *Th*_(*C*)_ include false positives due to the small sequential speckles (Fig. 2D, *upper*) or a combination of these and the one-frame appearance of a fluorophore (Fig. 2D, *lower*). The false positives are reflected in the histogram of the average *I* values within a trajectory (*I*_ave_) (Fig. 2E). When *Th*_(*C*)_ = 3, a lower *I*-value population occurs than that of the manual picking (Fig. 2E, *green*). When *Th*_(*C*)_ increases to 5, the low *I*-value population decreases, but remains (Fig. 2E, *orange*). The additional use of *Th*_(*I*)_ and *Th*_(*σ*)_ or further increase in *Th*_(*C*)_ to 7 shows approximately the same distribution as manual picking (Fig. 2E, *magenta and blue*), indicating effective removal of the false positives.

Furthermore, test analyses of dwell time and mean square displacement (MSD) were conducted using the data screened with the thresholds showing fewer failures on the *I*_ave_ histogram. Because the dye’s fluorescence was zero in the blinking events, the allowance of one dark frame within a continuous spot can prevent the tracking from an interruption in the middle (Fig. 2C, *asterisks*), as previously reported (Gebhardt *et al*., 2013). The dwell time histograms and the MSD plot of the screened fluorescent spots identified differ from those of manual picking; the dwell time and MSD value are underestimated (Fig. 2F). More details are found in Supplemental Text (II) and Figures S3–S6.

Hence, we developed a three-step screening method applying an additional threshold of *I*_ave_ (*Th*_(*Iave*)_ [a.u.]). First, all ROIs where the iterative solution converged were roughly screened with *Th*_(*I*)_ = 250, *Th*_(*σ*)_ = 1, *Th*_(*R*)_ = 1, and *Th*_(*C*)_ = 3. Some false positives remained after the first step, as shown in the histogram of *I* (Fig. 2G, *green*). Second, *Th*_(*Iave*)_ was set as 600 to exclude 99% of the false positives, while missing 10% of the short dwell trajectories. The shape of histogram *I* is almost the same as that of manual picking (Fig. 2G, *blue*), indicating that the overall population included almost all correct single molecules after screening with *Th*_(*Iave*)_. Finally, the remaining spots were rescreened with thresholds of *Th*_(*R*)_ = 2 and *Th*_(*C*)_ = 5 to conjugate the short trajectory fragments. In the final step, up to four consecutive dark frames in an existing continuous trajectory were allowed, whereas only one was allowed in the first step. The shape of histogram *I* did not change before and after the final step (Fig. 2G *red*), indicating no further failures. The dwell time histograms and the MSD plot of Nanog-EGFP in the nucleus using this protocol was almost the same as those of manual picking (Fig. 2F, *red*). Considering that the illumination intensity in the HILOM varies for each cell, unlike the TIRFM, we additionally developed a method to adaptively set *Th*_(*I*)_ and *Th*_(*Iave*)_ to each cell. Thus, we established an automatized protocol to obtain the same analysis results as manual picking. More details are described in Supplemental Text (III) and Figures S7–S8.

### Analysis of dissociation rate of Nanog or Oct4-EGFP on its target loci

First, we analysed the dissociation rate of Nanog- or Oct4-EGFP from its target loci, which is the reciprocal of the interaction time, and investigated its dependence on the differentiating condition. Time-courses of parameter *I* on a fixed ROI clearly show the appearance and disappearance of a fluorescent spot (Fig. 3A). While the appearance corresponds to the binding events of Nanog- or Oct4-EGFP, the disappearance includes alternative possibilities of dissociation and photobleaching (Figs. 1D and 3A, *red arrows*). The dissociation rate of Nanog- or Oct4-EGFP was expected to be smaller than the photobleaching rate of fluorescent proteins in SMT regarding the dissociation rate of other TFs (Chen *et al*., 2014; Xie *et al*., 2017) and the photobleaching rate of EGFP (Yu *et al*., 2006; Presman *et al*., 2017; Peterman *et al*., 1999), previously reported. A method to solve this problem has been previously developed because the dissociation rate is independent of the frame period, and the photobleaching is dependent; they can be mathematically distinguished by collecting the dwell time data at various frame periods (Gebhardt *et al*., 2013; Chen *et al*., 2014). Thus, we collected the dwell events at 50, 100, 150, 250, and 450 ms frame periods and extracted the frame period independent term, *i.e.,* the dissociation rate *k_off_*, of Nanog- and Oct4-EGFP in an mESC (supplemental text (IV) and Figs. S9–S12). The dwell events were distributed throughout the nucleus regardless of the frame period (Fig. 3B).

**Figure 3.**
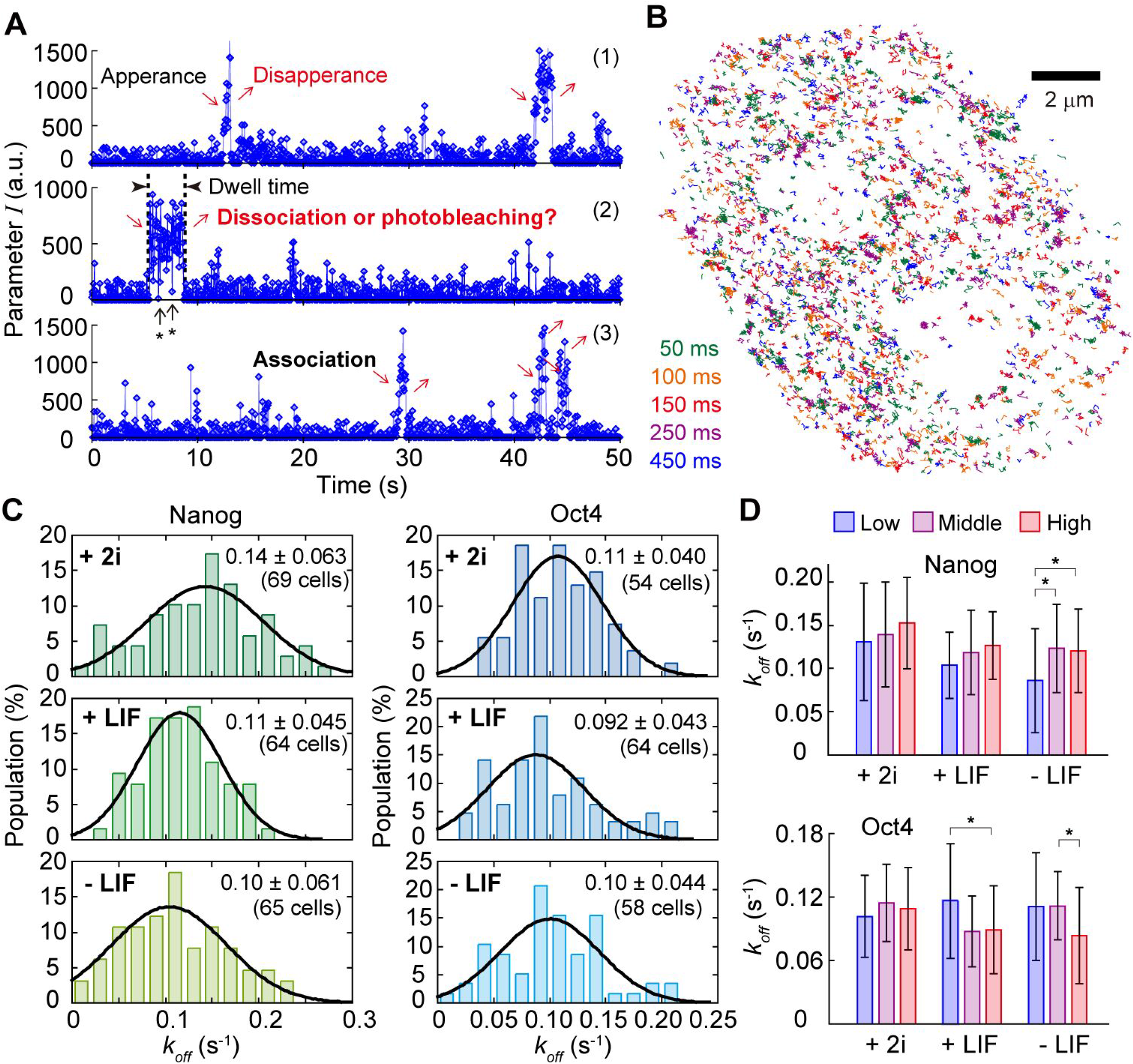
Analysis of dissociation rate of Nanog- or Oct4-EGFP on its target loci. (**A**) Typical three time-courses of parameter *I*, the fluorescent intensity of a single fluorophore, on a fixed ROI. Asterisks indicate blinking events of the fluorescent protein. Arrows indicate events of appearance and disappearance of the fluorescent spots. (**B**) A typical example of all dwell trajectories obtained within every 400 frames at frame periods of 50 (*green*), 100 (*orange*), 150 (*red*), 250 (*magenta*), and 450 ms (*blue*) in a nucleus in +LIF. (**C**) Histograms of dissociation rate of Nanog-EGFP (*left*) and Oct4-EGFP (*right*) in an mESC in +2i (*top*), +LIF (*middle*) and −LIF (*bottom*) conditions. Black line shows the fitting result with a normal distribution. (**D**) Dependence of expression level of Nanog-EGFP (*upper*) and Oct4-EGFP (*lower*) on dissociation rate. Considering the distribution of the fluorescent intensity (Fig. 1E), low (*blue*), middle (*magenta*), and high (*red*) were defined as cells showing the log value of the fluorescent intensity of < 1.9, 2.0–2.3, and > 2.4 [a.u.] for Nanog-EGFP or < 1.9, 2.0–2.2, and < 2.3 [a.u.] for Oct4-EGFP, respectively. Error bars are standard deviations. Asterisks indicate less than 0.05 of the p-value in the Student’s *t*-test.

The *k_off_* of Nanog- and Oct4-EGFP in an mESC exhibited different behavioural characteristics under the conditions of +2i, +LIF, and −LIF (Fig. 3C). The mean *k_off_* of Nanog-EGFP in +2i was 0.14 s^-1^, which significantly decreases to 0.11 s^-1^ in +LIF, and further decreases, but insignificantly to 0.10 s^-1^ in −LIF (Fig. 3C, *left*). Meanwhile, Oct4 exhibits no significant difference between the three conditions (Fig. 3C, *right*). By the classification into three categories depending on its expression level (*low*, *middle*, and *high*; see the legend of Fig. 3D), the mean *k_off_* positively correlates with the expression level in Nanog-EGFP, although it is statistically insignificant except for the low expression in–LIF (Fig. 3D, *upper*). However, almost no significant difference is observed between the expression levels of Oct4-EGFP; *k_off_* is negatively correlated in +LIF and −LIF (Fig. 3D, *lower*). Thus, Nanog-EGFP interacted longer with its target loci when differentiating and/or decreasing its expression level, whereas Oct4-EGFP did not.

### Analysis of fluctuating movements of Nanog- or Oct4-EGFP on its target loci

The chromatin condensation corresponding to the differentiation state reflects the behaviour of Nanog- or Oct4-EGFP on the locus and alters the physical properties of DNA. Next, the relationship between the differentiation state or the expression level and the physical properties of the binding site was investigated using the MSD analysis, thereby enabling the quantification of fluctuating movements (Fig. 4A) (Kusumi *et al.*, 1993; Saxton & Jacobson, 1997; Martin *et al*., 2002; Liu & Tjian, 2018). Because the average MSD curve appears saturated (Fig. 4B), the two parameters, confined radius (*R_c_*) and diffusion coefficient (*D*), could be estimated (Kusumi *et al.*, 1993; Miné-Hattab & Xavier, 2020). *R_c_* and *D* are thought to reflect chromatin condensation and local viscoelasticity, respectively, near the target loci of Nanog and Oct4 (Lionnet & Wu, 2021; Lerner *et al*., 2020; Miné-Hattab & Xavier, 2020).

**Figure 4.**
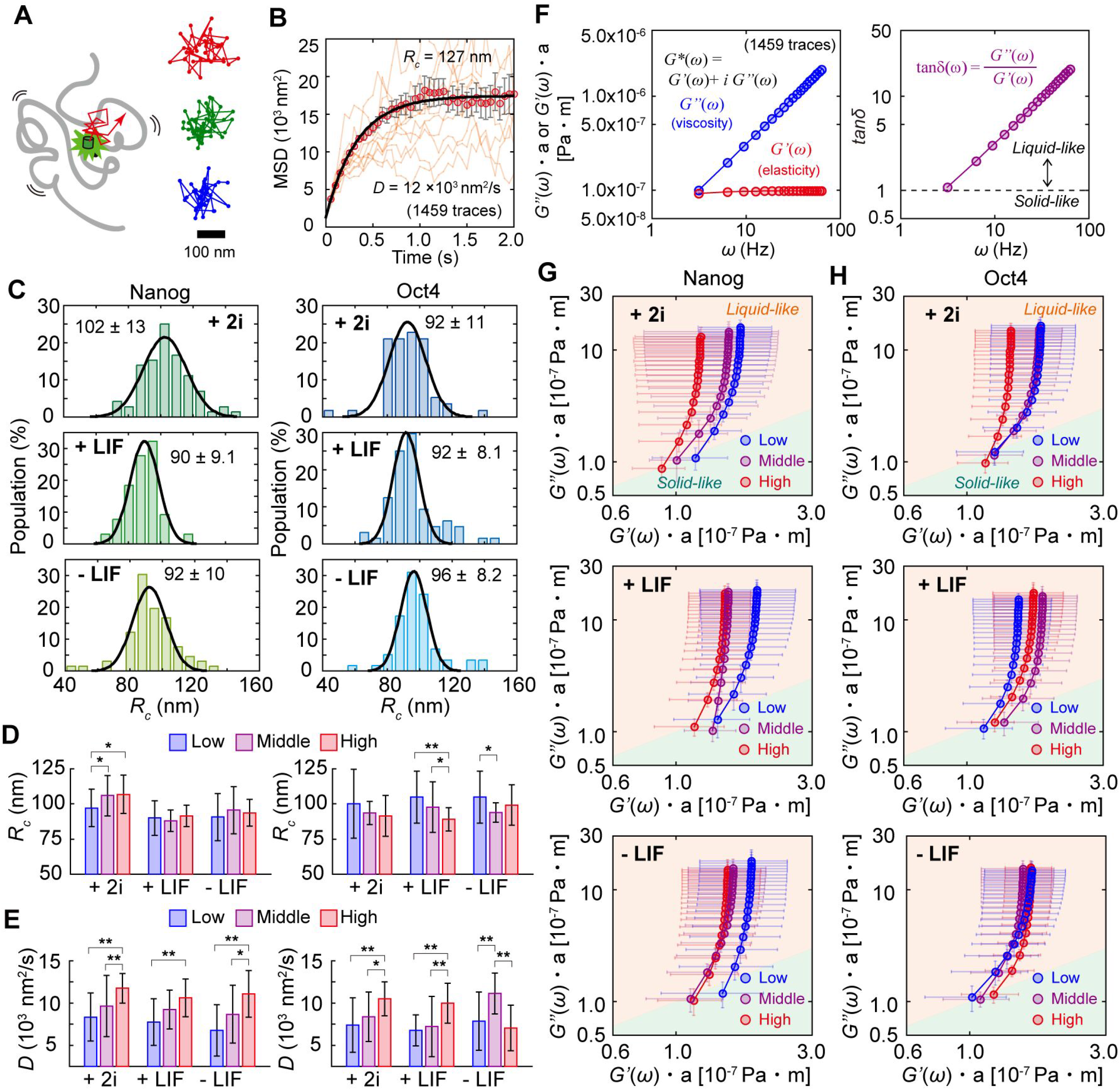
Analysis of fluctuating movements of Nanog or Oct4-EGFP on its target loci. (**A**) Cartoon of fluctuating movement of EGFP molecules on a DNA chain, and typical three trajectories of Nanog-EGFP. (**B**) A typical example of MSD plot of Nanog-EGFP in an mESC in +2i. The orange lines indicate each MSD curve obtained from a trajectory of an individual molecule. The red plots are the average of the single MSD curves within a cell. Error bars are the standard deviation. (**C**) Histograms of *R_c_* of Nanog-EGFP (*left*) and Oct4-EGFP (*right*) in an mESC in +2i (*top*), +LIF (*middle*), and -LIF (*bottom*). Black line shows the fitting result with a normal distribution. (**D, E**) Dependence of expression level of Nanog-EGFP (*left*) and Oct4-EGFP (*right*) on *R_c_* (**D**) or *D* (**E**). Asterisks and double asterisks indicate less than 0.05 and 0.01 of p-value in Student’s *t*-test, respectively. (**F**) A typical example of microrheology analysis of Nanog-EGFP in an mESC in +2i. Laplace transformation was applied to an average MSD (B, *red*). (G, H) *G”*(*ω*)-*G’*(*ω*) plot of Nanog-EGFP (**G**) and Oct4-EGFP (**H**) categorized with expression level. Error bars are standard deviations. The definition of low (*blue*), middle (*magenta*), and high (*red*) in **D**, **E, G**, and **H** were the same as that of Fig. 3D.

The *R_c_*-value of Nanog-EGFP significantly decreases after the removal of 2i, but does not further decrease after removing LIF (Fig. 4C, *left*), suggesting that the chromatin condensation near the Nanog’s target loci already began by removing 2i, which is consistent with the results of nuclear staining (Fig. 1F, *top panels*). Oct4-EGFP does not exhibit significant differences among the three conditions (Fig. 4C, *right*). In the expression dependence, the *R_c_* in +2i positively correlates with the Nanog-EGFP expression level, whereas those in +LIF and −LIF do not (Fig. 4D, *left*). The *R_c_* of Oct4-EGFP under all conditions tends to negatively correlate with the expression level (Fig. 4D, *right*), indicating that the DNA near the loci to which Oct4 bound was a flexible structure when mECSs expressed less Oct4. Overall, the *Rc* analysis results were similar to those of *k_off_* (Fig. 3CD), implying a relationship between the dissociation rate and chromatin condensation. Meanwhile, there was no significant difference between the *D*-value among the conditions in both Nanog- and Oct4-EGFP, and it negatively correlated with the expression level in both Nanog- and Oct4-EGFP, except for the −LIF condition in Oct4-EGFP (Fig. 4E).

To further consider viscoelasticity, the dynamic moduli of elasticity and viscosity, storage shear modulus (*G’*(*ω*)), and loss shear modulus (*G”*(*ω*)) were separately calculated since the generalized Stokes–Einstein relationship connects the MSD to the complex shear moduli in the Laplace domain (Ferry, 1980; Shinkai *et al*., 2020a; Shinkai *et al*., 2020b). *G”*(*ω*) linearly correlates with the frequency (*ω*) in all cases and is more than 10-fold > *G’*(*ω*) in all cases (Fig. 4F, *left*). The ratio of *G”*(*ω*) and *G’*(*ω*) is almost >1.0, indicating a liquid-like feature rather than a solid-like feature (Fig. 4F, *right*).

The *G’*(*ω*) of Nanog-EGFP matches the general expectation based on the chromatin condensation: higher expression or less differentiation, lower *G’*(*ω*); lower expression or more differentiation, higher *G’*(*ω*) (Fig. 4G). Generally, when *G’*(*ω*) and *G”*(*ω*) are proportional to *ω*^2^ and *ω*, respectively, the system relaxes for the slowest molecular motion. Here, only *G”*(*ω*) relaxes and shows a fluidic behaviour, and the elastic component behaviour of *G”*(*ω*) deviates from this trend. Furthermore, because the comparison with the differentiation state and the expression level exhibited a change in *G’*(*ω*), can be interpreted as the binding of Nanog promotes the elastic behaviour of DNA properties. Meanwhile, Oct4-EGFP showed interesting behaviour. While the tendency is almost the same as that of Nanog-EGFP in +2i (Fig. 4H, *top*), the lower expression decreases *G’*(*ω*), lowering the elasticity in +LIF (Fig. 4H, *middle*). The removal of LIF eliminates the expression dependence of the *G”*(*ω*)–*G’*(*ω*) plot (Fig. 4H, *bottom*). These results indicate that the Oct4-EGFP interaction weakens the DNA chain elasticity when the Oct4-EGFP expression decreases.

### Analysis of interaction clustering of Nanog or Oct4-EGFP on its target loci

Visualizing the dwelling frequency per unit area highlighted the clustering of the dwell events by removing 2i (Fig. 5A). It has been reported that TFs have different affinities against each locus and preferentially bind to high-affinity loci, corresponding to the cellular states (hotspots) observed under the light-sheet illumination (Liu *et al*., 2014; Kitagawa *et al*., 2017). We quantified the clustering of the dwell events, more specifically, and evaluated the size and distribution of hotspots of Nanog- or Oct4-EGFP during differentiation using a pair correlation analysis (Sengupta & Lippincott–Schwartz, 2012; Sengupta *et al*., 2013; Liu *et al*., 2014). The pair correlation function of all dwells (*g*(*r*)), where *r* is the distance, usually exhibits a single exponential curve, where the exponential decay constant (*ξ*) and the intercept (*g*_0_) reflect the mean cluster size and mean probability of the dwell events at a clustering site, respectively (Fig. 5B).

**Figure 5.**
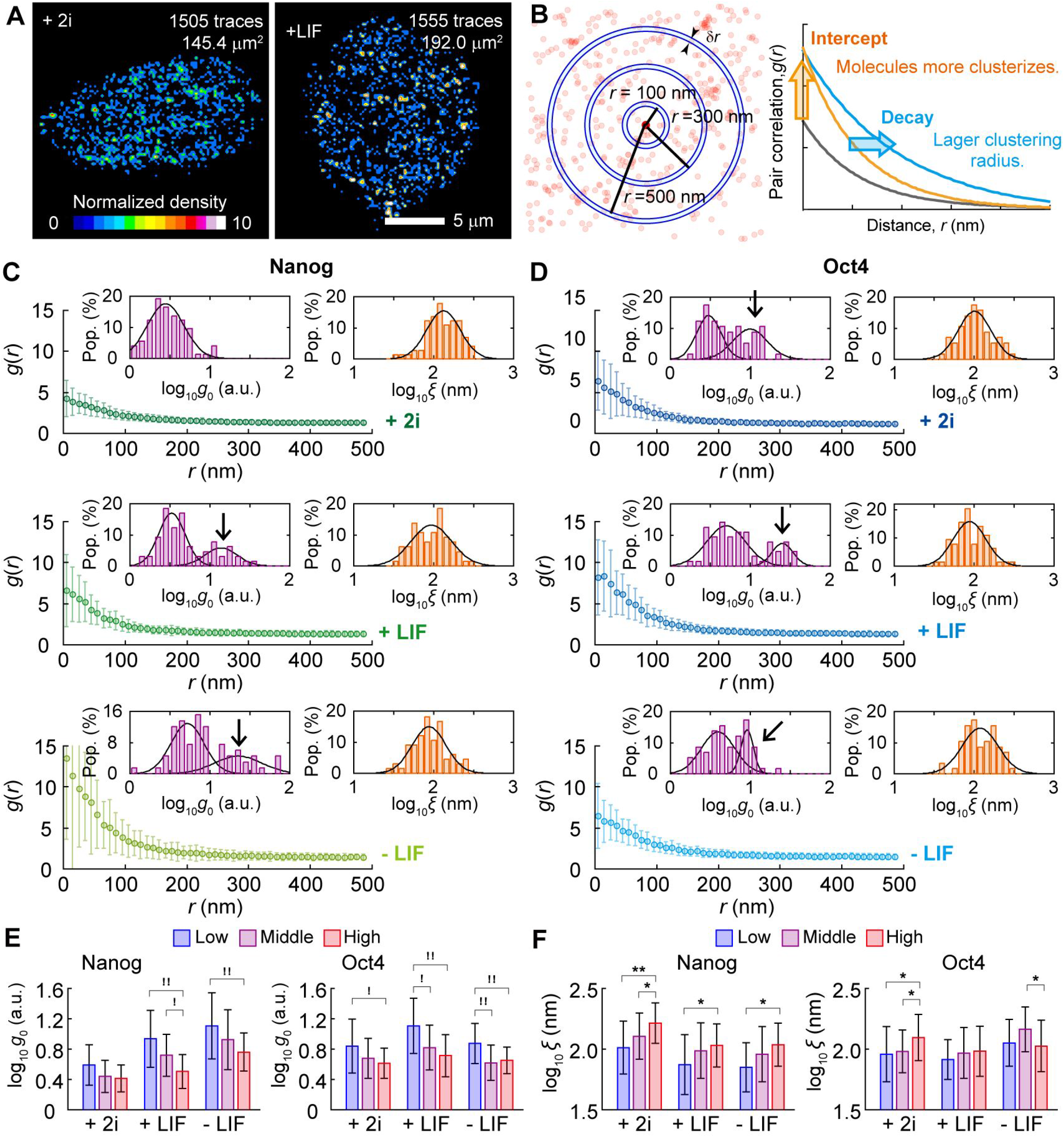
Analysis of clustering of dwell events of Nanog- or Oct4-EGFP on its target loci. (**A**) Visualization of the dwelling frequency of Nanog-EGFP per unit area, referred to as density, in an mECS in +2i (*left*) and +LIF (*right*). The colour corresponds to the density of Nanog dwell events (events/μm^2^) within 2,000 frames. (**B**) Explanatory drawing of the pair correlation analysis. The calculation of the probability of Nanog- or Oct4-EGFP molecule dwelling to another site at a given distance from a dwelling site provides the comparison to randomly dwelling. (**C, D**) The averaged correlation index *g*(*r*) of Nanog-EGFP (**C**) and Oct4-EGFP (**D**) in +2i (*top*), +LIF (*middle*) and -LIF (*bottom*). Insets are histograms of logarithm value of the intercept *g*_0_ (*magenta*) and the exponential decay constant *ξ* (*orange*) of single mESCs. Black lines in insets show the fitting results with a normal distribution. (**E, F**) Dependence of expression level of Nanog-EGFP (*left*) and Oct4-EGFP (*right*) on log_10_*g*_0_ (**E**) and log_10_ξ (**F**). Error bars are standard deviations. The definition of low (*blue*), middle (*magenta*), and high (*red*) in E and F were the same as that of Fig. 3D. Asterisks and double asterisks indicate less than 0.05 and 0.01 of p-value in Student’s *t*-test, respectively. Exclamation marks and double exclamation marks indicate less than 0.05 and 0.01 of p-value in Mann Whitney’s *U*-test. respectively.

The average trace of *g*(*r*) decreases in the cluster size and increases the probability of the dwell events as the differentiation progresses for both Nanog-EGFP and Oct4-EGFP (Fig. 5CD). The average *g*_0_ of the Nanog-EGFP clustering increases after removing 2i and further increases by removing LIF on average (Fig. 5C). The histograms of the logarithm of *g*_0_ exhibit the appearance of the second population with large values with the removal of 2i, and the mean values of the two populations increase with the removal of LIF (Fig. 5C, *inset, left*). The average value of *ξ* significantly decreases with the removal of 2i, and not of LIF (Fig. 5C, *inset, right*). The *g*(*r*) of Oct4-EGFP increases with the removal of 2i, as did Nanog-EGF, and recovers with the removal of LIF (Fig. 5D). Two populations are also observed even in the presence of 2i, and the mean value of the second population increases with the removal of 2i and markedly decreases with the removal of LIF (Fig. 5C, *inset, left*). The average value of *ξ* does not change by removing 2i, but significantly decreases by removing LIF (Fig. 5D, *inset, right*). The expression dependence of log_10_*g*_0_ exhibits a negative correlation under all conditions in both Nanog- and Oct4-EGFP (Fig. 5E). Meanwhile, those of log_10_ξ exhibit a negative correlation in almost all cases, except in the −LIF condition of Oct4-EGFP (Fig. 5F).

### Change in parameters two days after removal of 2i and LIF

Two days after removing 2i and LIF, the expression levels of both Nanog-EGFP and Oct4-EGFP decreased, making it challenging to simply compare the single-molecule data at the same expression level before the −LIF condition (Fig. 1E). Nevertheless, considering the importance of the data obtained in the differentiated mESCs, we collected the data in mESCs with residual fluorescence in the −LIF2d condition and performed the same analyses. The *k_off_* of Nanog-EGFP returned to the value in +2i (Fig. 6A, *upper*), whereas that of Oct4-EGFP exhibited a slight, but insignificant, increment (Fig. 6A, *lower*). The viscoelasticity obtained using the microrheology analysis shows no obvious change in Nanog-EGFP (Fig. 6B, *upper*). In Oct4-EGFP, *G’*(*ω*) remarkably increases in the case of low expression, indicating that the DNA transferred more elastically (Fig. 6B, *lower*).

**Figure 6.**
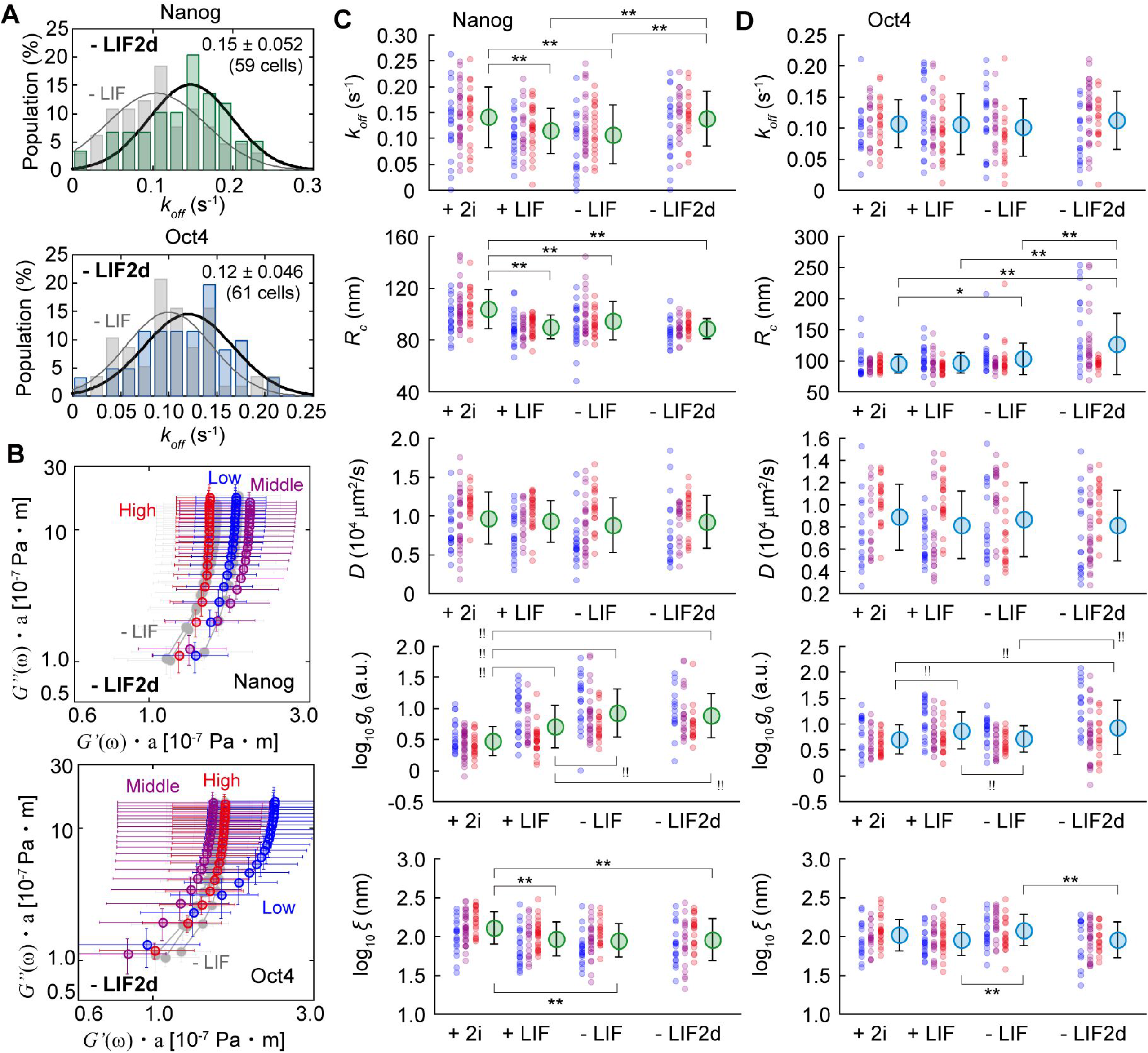
Data obtained 2 days after removal of 2i and LIF and summary of all data. (**A**) Histograms of dissociation rate of Nanog-EGFP (*upper*) and Oct4-EGFP (*lower*) in an mESC in the condition of -LIF2d. Black line shows the fitting result with a normal distribution. (**B**) G’’(ω)–G’(ω) plot of Nanog-EGFP (*upper*) and Oct4-EGFP (*lower*) categorized with expression level. In -LIF2d, the low (*blue*), middle (*magenta*), and high (*red*) were defined as cells showing the log value of the fluorescent intensity of < 1.7,1.8–2.1, and > 2.2 [a.u.] for Nanog-EGFP or < 1.5, 1.6–1.9, and > 2.0 for Oct4-EGFP, respectively. (**C, D**) Plots of all data of cells categorized with expression level (*red, magenta, and blue*) and the mean values (*green* or *cyan*) in each parameter in each condition of Nanog-EGFP (**C**) or Oct4-EGFP (**D**). Asterisks and double asterisks indicate less than 0.05 and 0.01 of p-value in Student’s *t*-test, respectively. Exclamation marks and double exclamation marks indicate less than 0.05 and 0.01 of p-value in Mann Whitney’s *U*-test, respectively. The definition of low (*blue*), middle (*magenta*), and high (*red*) in +2i, +LIF, and -LIF were the same as that of Fig. 3D, and those in -LIF2d were the same as B in this figure. Error bars are standard deviations. Note: the data obtained in -LIF2d could not simply compared with the others because the expression levels differed between them.

In Nanog-EGFP, the dependence of the obtained parameters on the differentiating conditions correlates with each other until a day after the removal of 2i and LIF (Fig. 6C). The additional day did not alter *R_c_, D, g*_0_, and *ξ*, except for *k_off_*, reducing the correlation (Fig. 6C, *-LIF2d*). The dependence on the expression level shows the same trend in all conditions and parameters (Fig. 6C, *red*, *blue,* and *magenta*). However, *k_off_* and *D* do not depend on the differentiating condition in Oct4-EGFP (Fig. 6D, *top* and *middle*). The *R_c_*-value in Oct4-EGFP increased by an additional day, especially in middle and low expressions (Fig. 6D, *second top*), and *g*_0_ and returned to the value in +LIF (Fig. 6D, *second bottom* and *bottom*). The dependencies on the expression level also returned to that in +LIF (Fig. 6C, *red*, *blue*, and *magenta*).

## Discussion

Herein, we quantified the dissociation rate, fluctuation, and interaction clustering of single molecules of Nanog and Oct4 in a living cell nucleus using SMT based on the HILOM. This study required numerous single-molecule data of Nanog- and Oct4-EGFP in mESCs. We collected approximately more than 500 videos (including those for confirmation of reproducibility and preliminary experiments) comprising 4,000 images, each including 500–2,000 trajectories of single molecules. It was unrealistic to analyse all fluorescent spots corresponding to single molecules in the collected 400 × 4,000 images visually. Unfortunately, the previous automation method of the single-molecule identification for TIRFM (Yasui *et al*., 2018) could not work well in single-molecule imaging using the HILOM (Fig. 2F). The three-step screening proposed herein realized automated identification of single molecules in the HILOM observation (Fig. 2G, S7, and S8, and Supplemental Text (III)). This concept and procedure of the three-step screening may apply to the other in-nucleus single-molecule combined with a fluorescent protein. Recently, an analysis framework has been developed that provides theoretically optimized parameter settings to exhibit an overall constant probability of tracking failures in the conventional protocol (Kuhn *et al*., 2021). Moreover, deep learning or Bayesian inference methods have been developed to automatize single-molecule identification (Xu *et al*., 2019; Smith *et al.*, 2019). These techniques are powerful candidates for parallel use with this method in the near future.

To the best of our knowledge, there are only a few reports quantifying the binding kinetics of core TFs in mESCs. Single-molecule dynamics of Sox2, which is the other core TF, have been investigated in detail, including its association with Oct4 (Chen *et al*., 2014). The intranuclear movement of TFs is divided into two broad categories: target site exploration and binding to the target site. Concerning Sox2, most populations (~97%) were the former with a short association time (*k_off_* = ~1.25 s^-1^), and only a few (~3%) were the latter with a long association time (*k_off_* = ~0.08 s^-1^). They also observed the *k_off_*-value of the long association time for Oct4 in a fibroblast cell line (~0.07 s^-1^). They reported the *k_off_* values of the other pluripotency-related TFs: ~0.12 and 0.10 s^-1^ for STAT3 and ESRRB (Xie *et al*., 2017). Although our method could only capture the slower lower, the obtained *k_off_* values of Nanog-EGFP and Oct4-EFGP are consistent with these previously reported values. Chen *et al.* also reported that the residence time of Sox2 was elongated by overexpressing Oct4 in Oct4-negative cells (Chen *et al*., 2014). Herein, an increase in *k_off_, i.e.*, a decrease in the residence time, was observed two days after the removal of 2i and LIF when the expression of Nanog and Oct4 was almost extinct (Fig. 6A). These results suggest that both Nanog and Oct4 dissociation would be promoted during differentiation based on the insight that the transcription activity would be downregulated with differentiation, as in the case of the previously reported Sox2. Thus, our measured *k_off_* values are considered plausible.

Interestingly, when Nanog or Oct4 expression is incompletely reduced at the onset of differentiation, the *k_off_*-value of Nanog-EGFP decreases by the removal of 2i, indicating that Nanog interacted longer at its target loci (Fig. 3C, *left*). We also observed that the *k_off_-value* of Nanog-EGFP positively correlates with the expression level of Nanog (Fig. 3D, *upper*). Paradoxically, it can be interpreted that the prolonged Nanog interaction time contributes to recovering the expression of the pluripotency-related transcription genes, including itself, and decreases by the removal of 2i and in a lower expression state, forming a positive feedback loop for hindering the escape from pluripotency to the undifferentiated state. This mechanism might disappear when the mESCs are at the later stage of differentiation, increasing the *k_off_*-value of Nanog-EGFP (Fig. 6A, *upper*). Meanwhile, that of Oct4-EGFP does not depend on the differentiating condition and negatively correlates with the expression level (Fig. 3C, *right* and D, *lower*). Thus, the results of the *k_off_* analysis indicate that Nanog works as a gatekeeper by controlling its interaction time.

The obtained values of *D* for Nanog- and Oct4-EGFP agree excellently with the result of Sox2 bound to chromatin in living mESCs (0.017–0.025 μm^2^/s) (Liu *et al*., 2014). According to a previous report, where the *D*-value of proteins bound to heterochromatin was smaller than that of euchromatin (Piccolo *et al*., 2019), the positive correlation of *D* to the expression level of both Nanog- and Oct4-EGFP (Fig. 4E) is thought to reflect the transition from euchromatin to heterochromatin with differentiation. Additionally, the obtained *R_c_*-value was consistent with that of histone H2B in mESCs (Gómez-García *et al*., 2021), and *R_c_* decreases depending on the differentiation condition, but not on the expression (Fig. 4CD, *left*), implying that *R_c_* reflects the chromatin condensation during differentiation. Meanwhile, the positive correlation of *D* and the expression level for Oct4-EGFP are distorted with the removal of LIF (Fig. 4E, *left*), and the *R_c_*-value of Oct4-EGFP does not depend on the differentiation condition and tends to negatively correlate with its expression (Fig. 4CD, *right*). These results implied that the local viscoelasticity of DNA basically becomes more elastic with the euchromatin–heterochromatin transition during differentiation, and only the local area of the target loci of Oct4-EGFP reverts to its original property. Hence, it is speculated that the chromatin near the loci where Oct4 interacts opened when its expression decreased. The *G”*(*ω*)-*G’*(*ω*) correlation plot of Nanog-EGFP in the microrheology analysis further visualizes the increase in the DNA elasticity near the Nanog’s target loci following differentiation (Fig. 4G). Meanwhile, the DNA on which Oct4-EGFP is bound is less elastic in the low expression state with the removal of 2i (Fig. 4H, *middle*), and the difference among expression levels disappears with the additional removal of LIF (Fig. 4H, *bottom*). The chromatin condensation seemed to reopen near the loci bound by Oct4-EGFP as if it resisted differentiation. Additionally, the *k_off_* of Oct4-EGFP is in synchronism with those (Fig. 3C, *right* and D, *lower*). Hence, the results for Oct4-EGFP can be explained by the role of Oct4 in recruiting remodelling factors to reopen the condensed chromatin.

So far, we have discussed the assumption that the diffusion motion couples with the state transition of the euchromatin–heterochromatin transition. However, we would like to try another interpretation; the formation of super-enhancer regions during the ESC differentiation. With the same analysis herein, the clustering of the dwell events of Sox2 has been observed in a LIF-treated mESC (Liu *et al*., 2014). They claimed that the Sox2 interaction cluster resulted from the local enrichment of enhancers to which Sox2 binds. Herein, the appearance of the interaction clusters is represented as the second population of the larger value in the *g*0 histogram (Fig. 5CD, *arrows*). Because Sox2 binds to DNA pre-emptively and recruits Oct4 to its binding site (Chen *et al*., 2014), the second population of Oct4 is probably associated with the interaction clusters, including Sox2 (Fig. 5D). The second population is mainly derived from the lower-expressing mESCs in both Nanog and Oct4 (Fig. 6CD, *second bottom*), demonstrating that the downregulation of Nanog- or Oct4-EGFP expression is strongly related to the appearance of the high-affinity interaction site, and that Nanog-EGFP was observed only after removal of 2i and grew after LIF removal (Fig. 5C). Considering the above results, the enhancer regions with which Nanog interacts are thought to assemble into an interaction cluster when escaping from the pluripotency state. Since the formation of a super-enhancer causes local chromatin condensations (Zhang *et al*., 2021), this interpretation does not contradict the MSD results. Alternatively, the decrease in the number of loci with which Nanog and Oct4 interact along with the euchromatin–heterochromatin transition relatively increases the affinity of the remaining loci to Nanog or Oct4, causing the emergence of interacting clusters. Thus, unfortunately, we cannot conclude here whether the origin of the chromatin condensation is the euchromatin–heterochromatin transition or the formation of a super-enhancer. Further experiments, such as dual SMT of super-enhancer region and TFs, are needed for clarification.

The mean values and dependency on the differentiation condition of *ξ* are almost the same as those of *R_c_* (Fig. 4D and 5F), and those on the expression level are similar to those on *D* (Fig. 4E and 5F). Hence, the *ξ*-value is not directly attributed to the size of an interaction cluster, but to the diffusion movement. We noted the correlation of the logarithm of *g*_0_ or *ξ* and *k_off_*. While the *g*_0_–*k_off_* correlation depends on the culture conditions, the *ξ*–*k_off_* correlation is interdependently distributed on the same linear relationship regardless of whether Nanog- or Oct4-EGFP and the culture conditions (Fig. 7A). This result implies that the dissociation rate of Nanog or Oct4 is inherently correlated with the mechanical property of DNA near its binding site, independent of the differentiation state, but not to interaction clustering. The relationship between the amount of Nanog expression with the chromatin condensation and the stiffness of the nucleus has been previously reported (Chalut *et al*., 2012). We also investigated the effect of the forcible chromatin decondensation by drug treatment with trichostatin A (TSA), a histone deacetylase inhibitor, on the *ξ*–*k_off_* correlation. The addition of TSA to the medium to chemically induce the decondensation of chromatin decreases *g*_0_ with or without LIF (Fig. 7B), reflecting the inhibition of the formation of interaction clusters. The *k_off_*-value of Nanog- and Oct4-EGFP returned to 0.12 ± 0.06 (s^-1^) and remained unchanged at 0.99 ± 0.04 (s^-1^), respectively, and the *ξ–k_off_* correlation disappeared, whereas the *ξ* value was unaffected (Fig. 7C). Therefore, *k_off_* is not deterministically regulated by the mechanical features, but is still related to interaction clustering.

**Figure 7.**
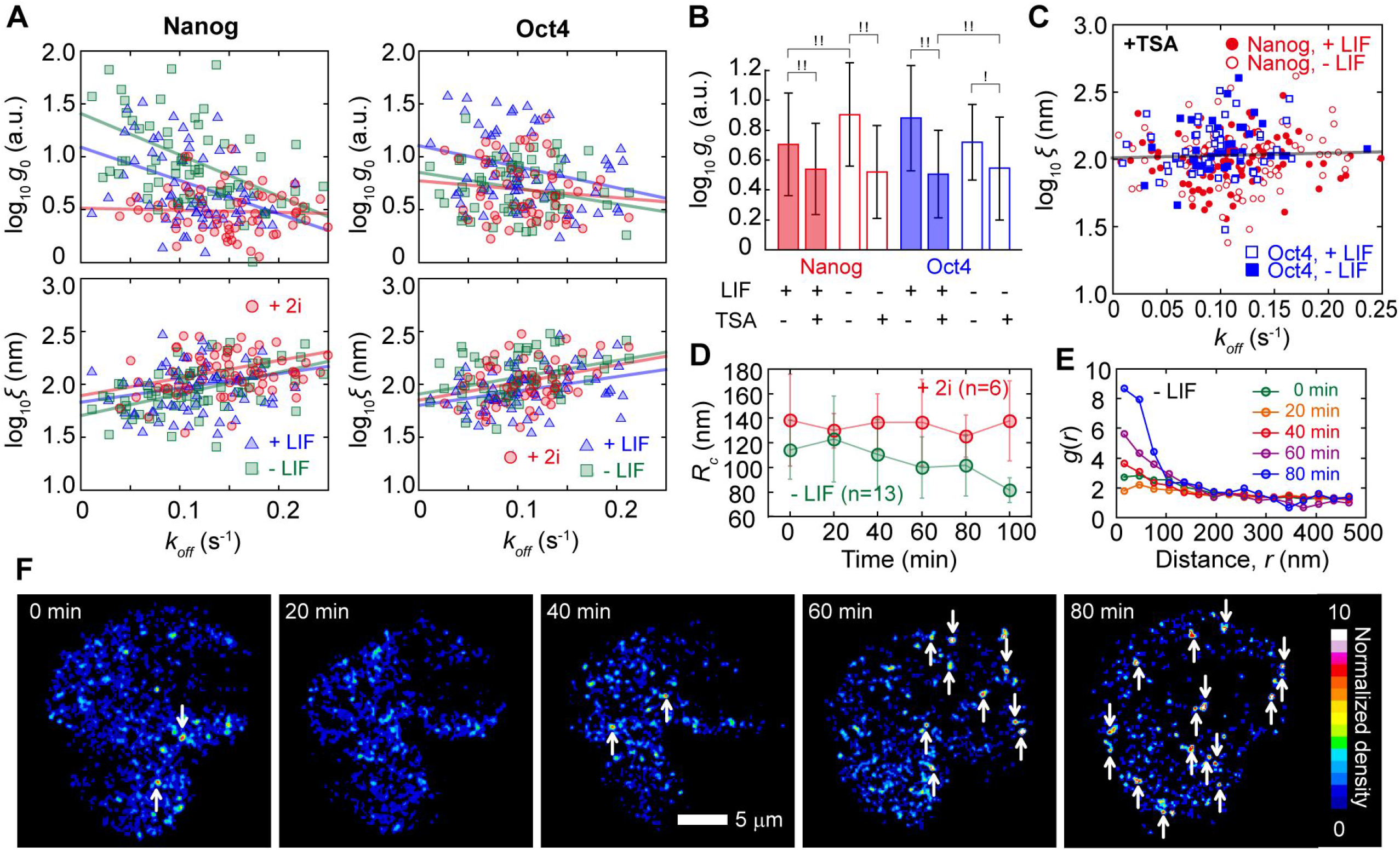
Additional analyses and experiments for the discussion. (**A**) Correlation plots of log_10_*g*_0_ (*upper*) or log_10_ξ (*lower*) and *k_off_* of Nanog-EGFP (*left*) and Oct4-EGFP (*right*) in 2i (*red*), +LIF (*blue*) and –LIF (*green*) conditions. Each plot indicates average value in a single cell. Lines are fitting results with a liner function. (**B**) Effect of TSA on log_10_*g*_0_ of Nanog-EGFP (*left*) and Oct4-EGFP (*right*) in +LIF (*filled*) and –LIF (*opened*). (**C**) Correlation plot of log_10_ξ and *k_off_* of Nanog-EGFP (*red circles*) and Oct4-EGFP (*blue rectangles*) in +LIF (*filled*) and –LIF (*opened*) with the addition of TSA. Plots are single cells. (**D**) Average time development of *R_c_* of Nanog-EGFP in 2i (*red*) and –LIF (*green*). (**E, F**) A typical example of time developments of correlation index *g*(*r*)(**E**) and visualization of the binding frequency of Nanog-EGFP per unit area (density)(**F**) in a mECS 6 hours after removal of 2i and LIF. Colour in F corresponds to the density of Nanog dwell events (events/μm^2^) within 2,000 frames. These were obtained from the same cell. Error bars in **A**, **B**, and **D** are standard deviations. Exclamation marks and double exclamation marks indicate less than 0.05 and 0.01 of p-value in Mann Whitney’s *U*-test, respectively.

Supposing that the chromatin condensation and the interaction clustering corresponded to the differentiation stage and the *R_c_* and *g*_0_ values for Nanog- and Oct4-EGFP reflect them, the changing moment of *R_c_* and *g*_0_ should be captured during the cell state transition. As expected, by observing the time development of Nanog-EGFP in the −LIF condition, we obtained data showing that the *R_c_* and *g*_0_ values dynamically increase within 100 min at any time after the LIF removal (Fig. 7DE). Using such data, the emergence of hotspots of functioning Nanog could also be visualized (Fig. 7F). While the strong photodamage had no or slight effect on the SMT, for example, the *R_c_* and *D* estimations in the surviving cells (Supplemental text VI and Figs. S15), the mESCs die from the critical photodamage during the long-term observation in most cases. Moreover, this event occurred stochastically 6 h after the LIF removal, making it difficult to collect repeated data. The automatization of the HILOM observation is needed to unveil the transfer of the chromatin state and the relationship to the binding behavioural characteristics of the TFs.

Two days after the removal of 2i and LIF, the previously decreased *k_off_*-value of Nanog-EGFP due to differentiation returned, and the DNA near the target loci of Oct4 became noticeably elastic during the low expression (Fig. 6). The parameter values of Oct4-EGFP returned to those before the LIF removal (Fig. 6D), implying that Oct4 stabilized the cellular state for a day after the LIF removal. However, only *R_c_* further increases (Fig. 6D, *second top*), and an interpretation of this behaviour remains unprovided. Oct4 may move onto other transcription networks after finishing its role as a pioneer factor. At least, it can be claimed that the mESCs are transferred from the pluripotency state to the other state on the second day after the removal of LIF, and that the results until a day after the removal of LIF (Figs. 3–5) unveiled the dynamic behavioural characteristics of Nanog- and Oct4-EGFP in the intermediate state transition.

By comprehensively considering these results, the following working hypothesis was proposed for the coworking of Nanog and Oct4 (Fig. 8). The interaction of Nanog or Oct4 with its loci is prolonged as differentiation occurs, correlating to the degree of the chromatin condensation during differentiation. The chromatin modification for downregulation of Nanog or Oct4 expression or formation of super-enhancer regions results in a longer-term interaction, increasing the transcription efficiency of related genes, including themselves. However, the Oct4 interaction recruits remodelling factors on the binding site as a pioneer factor, which opens the closed-structured chromatin. This mechanism is a negative feedback circuit, causing a stable attractor for the pluripotent state. Then, dissociation might be promoted during differentiation progression to the point of no return. In fact, in mESCs with slight residual fluorescence two days after the removal of 2i and LIF, the *k_off_* values of both Nanog- and Oct4-EGFP increased, and the DNA became more elastic, indicating the chromatin condensation progression (Fig. 6). According to this hypothesis, since Oct4 contributes to opening its target site, the overall expression level would be stable. Meanwhile, since the opening of Nanog’s target site is Oct4-dependent, the expression level fluctuates, causing heterogeneous expression. This trend is consistent with a previous report that the expression of Nanog is more heterogeneous than that of Oct4 (Chambers *et al*., 2007). This negative feedback system might be a source of heterogeneity in Nanog expression.

**Figure 8.**
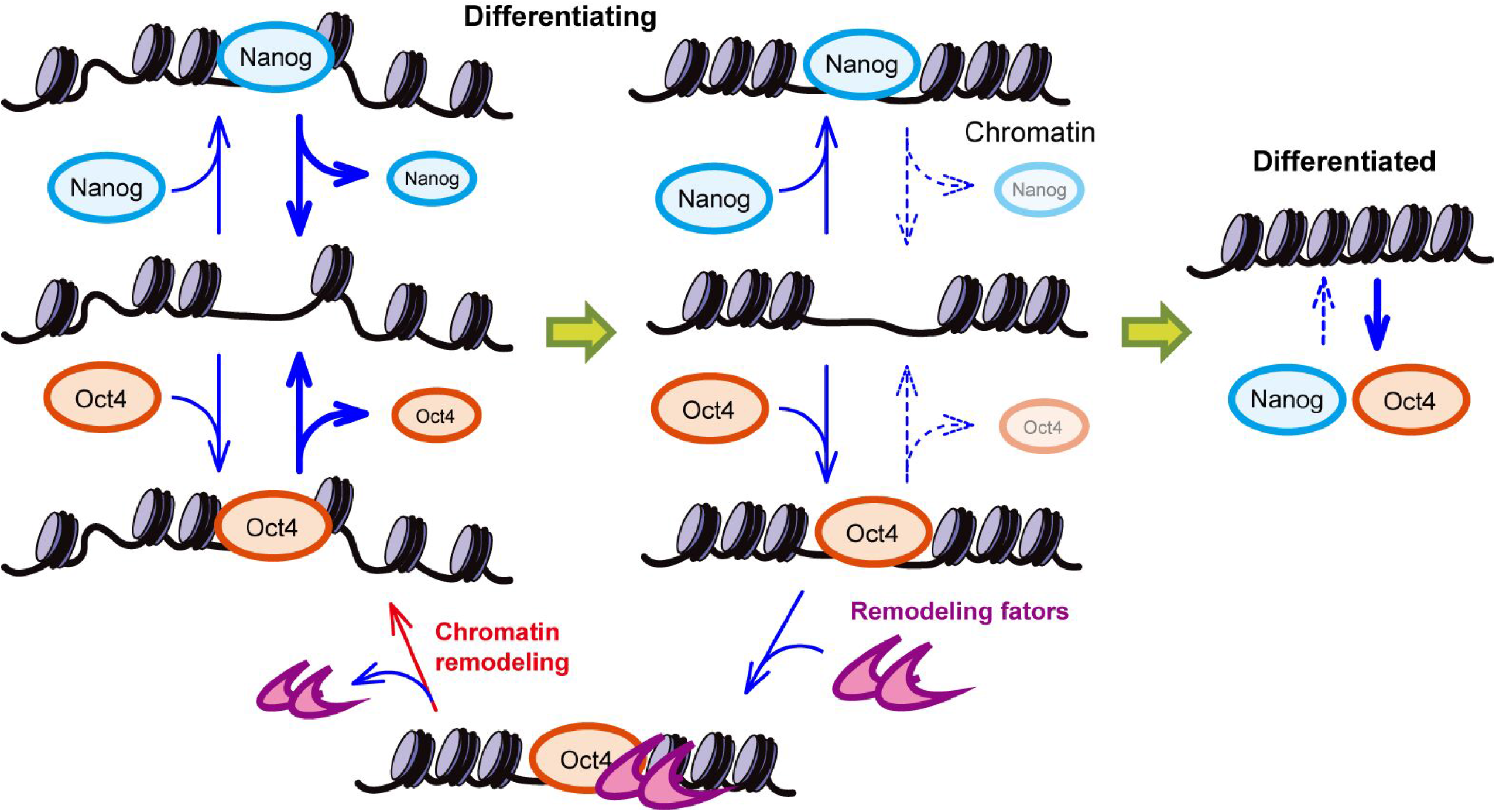
Proposed working model based on the results in this study. Nanog and Oct4 are indicated by blue ellipse and orange ellipse, respectively. The left is more undifferentiated, while the right is more differentiated. Chromatin is assumed to be condensed over time if without remodelling by pioneer factors. The interaction time of Nanog or Oct4 to its target loci positively correlated the degree of chromatin condensation. Once chromatin is completely condensed, Nanog or Oct4 cannot interact to the loci. The localization of Oct4 onto DNA recruits remodelling factors (*magenta*) to open condensing chromatin.

In conclusion, the SMT using the HILOM with EGFP enabled quantifying the on-site behavioural characteristics of Nanog and Oct4 in a living nucleus of mESC, biophysically supporting the current working model. Additionally, these quantitative data provided a new feedback mechanism for pluripotency maintenance, possibly a source of expression heterogeneity. Further studies are needed to confirm our hypothesis and investigate the causal relationship between the chromatin state and the interaction of Nanog or Oct4 with its target loci. Currently, we are preparing for an on-site simultaneous observation of Oct4-EGFP and chromatin opening at a single-molecular level (Supplemental Text VII and Fig. S14). Photodamage is the most significant hindrance to long-term observation. Also, more revolutionary innovations for the SMT are needed to fully elucidate the cofunctioning mechanism of Nanog and Oct4 in pluripotency maintenance.

## Materials and Methods

### Gene construction and cell line establishment of Nanog- or Oct4-GFP expressing mESCs

Herein, an EGFP was conjugated at the end of Nanog in only one allele using the transcription activator-like effector nucleases (TALEN) technique in an mESC (Hisano *et al*., 2013; Ota *et al*., 2013; Sakuma *et al*., 2013), E14tg2a cell line (AES0135, Riken Cell Bank, JP). The vector construction was based on the Golden Gate ligation (Sakuma *et al*., 2013). The donor plasmid was constructed using pBluescript II as the backbone and by ligating the insert DNA, comprising of the PGK promoter, puromycin R gene, and EGFP amplified using PCR. A pair of TALENs was designed in the intron before the last coding exon of Nanog. The target sequences of these TALENs were 5’-TGCCTGCCTAGTCTCAGGAGTGCTGGGGTTAACGGCCTGTGCGGCCA-3. The last coding exon in the genome was replaced by a sequence comprising the last coding exon fused with EGFP by co-transfecting a pair of TALEN vectors and a homologous donor. Transfected mESCs were cultured for two days, and the EGFP-positive cells were isolated using flow cytometry (BD FACS Aria III™, BD Biosciences, USA). The single isolated cells were expanded and used for the experiments. The Oct4-EGFP knock-in mESC line was established from blastocysts of Oct4-EGFP knock-in mice (RBRC06037, Riken Cell Bank) (Toyooka *et al*., 2008). The cell line was derived with some minor modifications, as previously described (Czechanski *et al.*, 2014).

The mESCs were seeded and cultured until compact colonies were formed. The colonies were molecularly confirmed via several stem cell markers, and the cells were expanded for use. All culture incubations were performed at 37°C, 5% CO_2_, and 95% humidity.

### Preparation of mESCs for single-molecule observations

Nanog-EGFP or Oct4-EGFP mESC lines were maintained in Dulbecco’s modified Eagle’s medium (DMEM; D6046, Sigma-Aldrich, USA) containing 10% FBS (16141–075, Gibco, USA), 1% penicillin-streptomycin (P4333, Sigma-Aldrich), 1% GlutaMAX-1 (35050–001, Gibco), 1% nonessential amino acids (11140–050, Gibco), 1% nucleosides (ES-008-D, Millipore, USA), 1% sodium pyruvate (S8636, Sigma-Aldrich), 0.1% 2-mercaptoethanol (60-24-2, Sigma-Aldrich), and 0.1% leukaemia inhibitory factor (LIF; NU0013–1, Nacalai, Japan) on 10-cm dishes (353003, BD Biosciences) coated with 0.1% gelatine (EmbryoMaxR 0.1% gelatin; ES-006-B, Merck Millipore, Germany). Both Nanog-EGFP and Oct4-EGFP expressing cells were passaged every two days.

Herein, the differentiating conditions were defined as follows: the presence of both LIF and 2i (hereinafter +2i) as the initial condition, one day after removing 2i (hereinafter +LIF), further removal of LIF for one day (hereinafter −LIF), and after one more day (hereinafter −LIF2d). For the +2i condition, the cells were cultured in the above culture medium containing the two inhibitors: 1 μM MAPK/ERK inhibitor (PD0325901; Stemolecule™ 04–0006, Stemgent, USA) and 3 μM GSK3 inhibitors (CHIR 99021; 04–0004, Stemgent, Stemolecule™). Twenty-four hours before microscopy, 1 × 10^5^ cells were seeded on 35-mm glass-bottom dishes (D11130H, Matsunami-Glass, Japan) coated with fibronectin (354008, Corning, USA) as per the commercial procedure under the 2i condition. Before microscopy, the medium was changed to phenol red (-) FluroBrite™ DMEM (A18967-01, Gibco). For the +LIF condition, 2i was removed when replating on the glass-bottom dish. For −LIF, mESCs were cultured in a medium containing LIF, but in the absence of 2i for 24 h, and replated onto the glass-bottom dish in the absence of 2i or LIF. For the trichostatin A (TSA) treatment, TSA was added at 0.5 μM in the culture medium when seeded on a glass-bottom dish 12 h before microscopy.

### Immunostaining

Cells were seeded as described above and fixed with 4% paraformaldehyde 12 h after seeding. After fixation, the cells were permeabilized with 0.5% Triton X-100 diluted in PBS for 10 min at room temperature (RT), blocked for 30 min at RT with CAS-Block^TM^ Histochemical Reagent (008120, Thermo Fisher Scientific), and incubated for an hour at RT with primary antibody against H3K27ac (1:2,000, ab4729, Abcam). The nuclei were stained by incubating the cells with PBS containing fluorescent secondary antibodies (1:1,000, A21207, Invitrogen) and DRAQ5™ (1:1,000, DR50200, BioStatus Limited) for an hour at RT. The stained cells were imaged using a confocal fluorescence microscope (FV3000, Olympus, Japan) with 488 nm/561 nm/640 nm lasers and a UPLXAPO40XO (N/A 1.40) objective lens.

### Single-molecule observations in a living nucleus

A commercial microscope system (N-STORM, Nikon, Japan) with an objective lens (CFI Apochromat TIRF 100XC Oil, Nikon, Japan) was used. The intermediate magnification was ×1.6 and the total magnification was ×160. According to previous literature, only the incident optical pathway in the microscope was customized for the HILOM (Tokunaga *et al*., 2008). The cells were incubated at 37°C, 5% CO_2_, and 95% humidity in a stage-top incubator (INU-TIZ-F1, Tokai hit, Japan). The sheet-formed laser was illuminated 2–3 μm above the glass surface. An electron-multiplying charge-coupled device camera (iXon3 893, Andor, UK) was used for image acquisition. The active pixels were set to 256 × 256 pixels, and the exposure time was fixed at 50 ms with the frame transfer mode. One pixel corresponded to 100 nm in the microscope. The back-illuminated sensor in the camera was cooled to −90°C using water circulation from a Peltier cooler.

The image acquisition scheme is explained as follows: a cell was identified via bright-field observation without laser illumination, and an epi-fluorescent image was acquired to monitor the total expression of Nanog-EGFP or Oct4-EGFP (Fig. 1C). Next, the illumination was changed from the epi-illumination to the HILO illumination. After inducing the photobleaching of almost all EGFP molecules within the illuminated area, image acquisition was initiated. We obtained five sets of 400 sequential images with various time intervals in the following order: 400, 0, 200, 50, and 100 ms (total of 2000 images) for the dwell time analysis, and 2,000 images with 0 ms intervals for the others, with 50 ms exposure time in a dataset. The total image acquisition time was ~10 min. This procedure was automated using built-in software in the microscope system. After video acquisition, we confirmed that the shape of the cells did not change. The data of mESCs that changed their shape were removed from the collection. All images were processed by the Gaussian blur with kernel size 3 × 3 to reduce ‘salt and pepper’ noise before SMT.

In the HILOM observation, the lens effect and scattering due to intracellular microstructures and/or vesicles disordered the shape of the sheet-formed light depending on the cell arrangements within the field of view. Therefore, searching for an mESC suitable for the HILOM observation was time-consuming. Considering the cell condition, the microscopic observation time in a dish was limited to 1 h. Additionally, to precisely define the differentiation state 24 h after LIF removal, the total observation time was limited to 2 h. Thus, we collected a maximum of four to five videos per day in two to three dishes. Hence, to collect 20 cell videos, four or five individual sample preparations on different days were needed.

The movements of single molecules of Nanog- or Oct4-EGFP were digitally recorded. All analyses described below were performed using homemade software programmed by C++ (Visual Studio 2008, Microsoft, USA) or Python3, and visualization of quantified data was conducted using the OpenCV library (ver. 2.4.3). The analyses were performed using a blind method; the information about the data, such as the buffer condition, was hidden during the analysis from the analyst until the results were obtained.

### Collection of single-molecule data in-nucleus

The following equation was used for the Gaussian fitting:

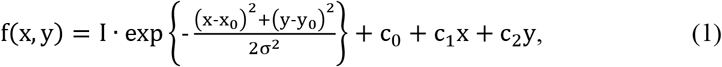

where, *c*_0_, *c*_1_, and *c*_2_ are the background fluorescence, assuming that the background signal inside the ROI is approximated by a tilted plane owing to unfocused fluorescence (Fig. 2A, *inset*) (Ichimura *et al*., 2014). The fitting computation was performed by an iterative calculation based on the Levenberg–Marquardt method (Levenberg, 1944) programmed with reference to “Numerical Recipes, 2^nd^ Edition,” (Chapter 15) (Press *et al*., 1992). All initial parameters for the Levenberg–Marquardt method were obtained automatically as follows (Ichimura *et al*., 2014): the initial parameters of *c*_0_, *c*_1_, and *c*_2_, were calculated with the linear least-squares method using only the outer boundary of the ROI. Because the logarithm of the Gaussian function is a simple quadratic function, the initial parameters of *x*_0_ and *y*_0_ were obtained using the linear least-squares method. The initial parameter of *I* is the pixel intensity at the initial (*x*_0_, *y*_0_) minus the background intensity calculated from *c*_0_, *c*_1_, and *c*_2_. The initial parameter of *σ* is simply a quarter of the ROI width. Afterward, the loop iterations of the Levenberg–Marquardt method were started. The ROI was scanned in steps of half the ROI diameter within a nucleus region, and the iterative calculation for the Gaussian fitting was performed at each step. The nuclear region was identified using epi-fluorescence images of Nanog- or Oct4-EGFP. Thus, all converging iterative solutions within the entire nucleus area were recorded into a collection for each mESC. The collection included the converged solutions even in the absence of fluorescent spots, which are false positives. The false positives were automatically removed as described in Supplemental Texts (II) and (III) and Figs. S3–S8.

### Dwell time analysis

We modified a previously developed method to reduce the photobleaching effect of the dwell time analysis (Gebhardt *et al*., 2013; Chen *et al*., 2014). The dwell time histograms from appearance to disappearance exhibited double exponential decay in all cases of 50, 100, 150, 250, and 450 ms frame periods. We globally fitted the histograms with the following equation:

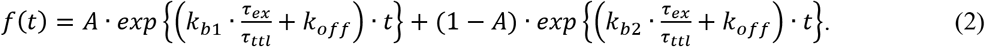

The details are described in Supplemental Text (IV) and Figures S9-S12.

### MSD analysis

The MSD analysis data were collected from 2,000 images acquired at a 50 ms frame period (20-Hz frame rate). Only trajectories comprising more than 10 positional data points were used. Expressing the position of a certain time as *f*(*t*), the MSD was calculated as follows:

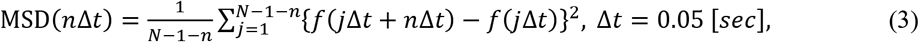

where, Δ*t* represents the frame period, *n* is the frame number, and *N* is the total number of single-molecule trajectories (Kusumi *et al*., 1993). The calculated MSD should include an estimation error in the position determination using Gaussian fitting. Therefore, the MSD obtained is as follows, as previously reported (Martin *et al*., 2002):

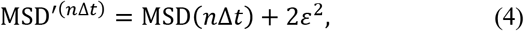

where, ε represents the estimation error. Because the calculated MSD exhibited confined diffusion, the MSD was approximated using the following equation:

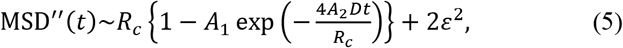

where *R_c_* and *D* are the plateau values of the MSD curve and diffusion coefficient, respectively. Also, *A*_1_ and *A*_2_ are constants determined by the confined geometry (Saxton & Jacobson, 1997). For simplicity, we fitted the MSD trace using the following equation by assuming symmetric diffusion (Miné-Hattab & Xavier, 2020):

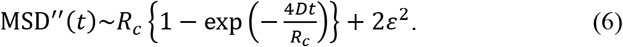

### Microrheology analysis for MSD data

To reduce the estimation error in the position determination*ε*^2^ [nm^2^], of each MSD data with the discrete-time *t_n_* = *n*·Δ*t* (*n* = 0, 1, –, 40 and Δ*t* = 0.05 [s]), we fitted with the first three time points (*n* = 1, 2, and 3) were fitted by the linear model, *f*(*t*) = a·*t* + 2·*ε*^2^ Next, each error-reduced set of MSD data were fitted using the following multi-component model:

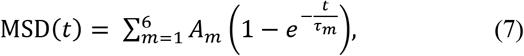

Here, the relaxation time (*τ_m_*) was set equal to 2 m^-1^·Δ*t*. To fit the values of 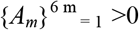, a SciPy function (scipy.optimize.curve_fit) was used with the trust region reflective method. Next, the MSD function was converted to dynamic compliance (Ferry, 1980; Shinkai *et al*., 2020a; Shinkai *et al*., 2020b) as a function of the frequency (*ω*) through the Fourier–Laplace transformation:

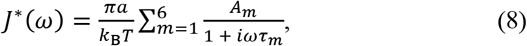

where *k*_B_ is the Boltzmann constant, *T* is the temperature, and *a* represents the radius of the probe particle. To calculate the complex modulus (*G*\*(*ω*)), the following relationship was used:

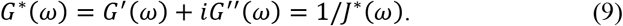

Since we were unable to determine the radius (*α*) in our experiments, we calculated the normalized storage and loss moduli (*G’*(*ω*)·*a* and *G*”(*ω*)·*a* [Pa·m] at *T* = 310 K. Since the discrete-time of the MSD data was *t_n_* = *n*·Δ*t* (*n* = 0, 1, …, 40), the discrete frequency (*ω_k_* = 2*π_k_* / 2.0 (rad/s)) (*k* = 1, 2, —, 20) was used.

### Analysis of clustering of dwell events

Before the clustering analysis, the mean centre position was calculated within a dwell trajectory as the dwell event position. Also, we calculated the probability of the existence of dwell events within a concentric tori of width (δ*r*) at a given distance (*r*) from the provided binding site (Sengupta & Lippincott-Schwartz, 2012; Sengupta *et al*., 2013). Because the estimation error in the position determination was reduced by averaging within a trajectory of <10 nm, the pair correlation distribution (*g*(*r*)) was approximated as follows:

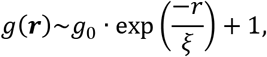

where and *g*_0_ provide rough estimations of the clustering radius and the average number of dwell events in a cluster, respectively (Sengupta & Lippincott-Schwartz, 2012).

## Acknowledgements

We thank Keiko Yoshizawa (RIKEN) and Kohei Yamamura (Osaka University) for preparation of the genetic construct, and Taishi Kakizuka (Osaka University) for investigation of correlation of immunofluorescence and EGFP fluorescence of Nanog-EGFP expressing mESCs. We would like to thank enago (www.enago.jp) for English language editing.

This work was supported by the Ministry of Education, Culture, Sports, Science and Technology [Grant-in-Aid for Scientific Research on Innovative Areas: JP18H05409 to T.M.W, JP18H05412 to S.O., JP16H06280, JP19H05794, JP19H05795 to Y.O. and JP20H05550 to S.O.], the Japan Society for the Promotion of Science [Kakenhi: JP19K16158 to K.O., JP21H03599 to T.M.W and JP19H03394 to Y.O.], and Japan Science and Technology Agency [CREST: JPMJCR1852 to T.M.W., JPMJCR15G2, JPMJCR20E2, and JPMJCR15G2 to Y.O; Moonshot R&D: JPMJMS2025-14 to Y.O.], and partially supported by the Japan Agency for Medical Research and Development [17bm0804008 to T.M.W.].

## Author contributions

K.O. performed all experiments and analyses, H.F. wrote the manuscript, S.S and S.O contributed to the microrheology analysis, Y.O. provided the TALEN technique, designed the experiments, and the critical discussion; K.A. contributed to establishing the mECS lines; and T.M.W. supervised the project and wrote the manuscript.

## Data availability

The single molecule videos used in the main results are available on the following site, https://ssbd.riken.jp/repository/193/, doi: 10.24631/ssbd.repos.2021.09.001. The other videos and analyses results can be made available from the corresponding author upon reasonable request.

